# Transparent transfer-free multilayer graphene microelectrodes enable high quality recordings in brain slices

**DOI:** 10.1101/2025.06.13.657566

**Authors:** Nerea Alvarez de Eulate, Christos Pavlou, Gonzalo León González, Lukas Holzapfel, Zhenyu Gao, Sten Vollebregt, Vasiliki Giagka

**Affiliations:** Department of Microelectronics, Faculty of Electrical Engineering, Mathematics and Computer Science, Delft University of Technology, Delft, The Netherlands; Department of System Integration and Interconnection Technologies, Fraunhofer Institute for Reliability and Microintegration IZM, Berlin, Germany; Department of Neuroscience, Erasmus MC, Westzeedijk 353, 3015 AA, Rotterdam, the Netherlands

**Keywords:** in vitro neural interfaces, electrophysiology, microelectrode arrays, graphene, multimodal applications

## Abstract

Resolving the underlying mechanisms of complex brain functions and associated disorders remains a major challenge in neuroscience, largely due to the difficulty in mapping large-scale neural network dynamics with high temporal and spatial resolution. Multimodal neural platforms that integrate optical and electrical modalities offer a promising approach that surpasses resolution limits. Over the last decade, transparent graphene microelectrodes have been proposed as highly suitable multimodal neural interfaces. However, their fabrication commonly relies on the manual transfer process of pre-grown graphene sheets which introduces reliability and scalability issues. In this study, multilayer graphene microelectrode arrays (MEAs) with electrode sizes as small as 10-50 µm in diameter, are fabricated using a transfer-free process on a transparent substrate for in vitro multimodal platforms. Through acute experiments using cerebellar brain slices, their ability to detect spontaneous extracellular spiking activity from neural cells, with a high signal-to-noise ratio up to 30-40 dB, is demonstrated. The recorded signal quality is found to be more limited by the electrode-tissue coupling than the MEA technology itself. Overall, this study shows the potential of transfer-free multilayer graphene MEAs to interface with neural tissue, which paves the way to advance neuroscientific research through the next-generation of multimodal neural interfaces.

## 1. Introduction

The human brain is one of the most complex biological systems, comprising billions of neurons that communicate through intricate neural networks. These networks are the basis of essential functions, such as cognition, memory, and behavior, as well as neurological disorders such as epilepsy, Parkinson’s disease, and schizophrenia [1], [2], [3]. Unraveling the mechanisms of these networks is a fundamental challenge in neuroscience, requiring advanced tools capable of mapping neural activity with high spatial and temporal resolution [4].

Neural interfaces have emerged as powerful tools for interacting with the nervous system, enabling the detection and modulation of electrical signals from excitable cells [5]. Conventional electrophysiological techniques, such as patch-clamp recordings and metal-based microelectrode arrays (MEAs), have been the gold standard for studying neuronal activity and have significantly advanced our understanding of neural circuits [6], [7]. However, these technologies are limited in their ability to capture the dynamics of large-scale neural networks across extended fields of view, hindering progress in understanding complex brain functions.

Recently, there has been a growing interest in optical techniques in neuroscience due to their ability to record and manipulate neural activity with increased spatial resolution. Methods such as calcium imaging and optogenetics allow researchers to simultaneously monitor the activity of hundreds to thousands of neurons, providing insights into neural connectivity and network dynamics [8], [9], [10]. Despite their advantages, optical techniques alone often lack the temporal resolution required to capture fast electrophysiological events, such as action potentials, with millisecond precision. Multimodal neural platforms that integrate optical and electrical modalities offer the potential to further advance the depth of understanding of our nervous system by offering greater spatiotemporal resolution [11], [12].

Graphene has emerged as a highly attractive material for such neural interfaces owing to its unique combination of biophysical, electrical, mechanical, biological, and optical properties [13], [14], [15], [16], [17]. This 2D material exhibits flexibility and mechanical resilience, which allows it to be integrated into highly conformable soft interfaces, conforming closely to the complex geometries of neural tissues and reducing the mechanical mismatch. Furthermore, the biocompatibility of graphene, which was initially controversial, has shown promising results in more recent studies. In particular, graphene produced by chemical vapor deposition (CVD), shows no significant signs of toxicity, supporting its use in chronic neural interfacing applications [18]. Due to its intrinsic optical transparency, monolayer CVD graphene has been explored for simultaneous electrophysiological recordings and optical imaging [19]. However, the inherent quantum capacitance of graphene poses a challenge when scaling down electrode sizes, as it limits the impedance of the electrodes. To address this limitation, researchers have explored various strategies, including the chemical doping of graphene [20] and surface modifications based on the integration of platinum nanoparticles [14], [16]. These approaches have successfully achieved lower impedance and higher charge storage capacity (CSC) values, enhancing the performance of graphene-based neural interfaces.

Despite their potential, the widespread adoption of graphene-based neural interfaces has been hindered by challenges in fabrication, particularly the need to transfer pre-grown graphene layers onto target substrates. This transfer process often introduces defects, contamination, and scalability issues, limiting the reliability and performance of graphene electrodes [21], [22]. CVD graphene electrodes have been developed using a transfer-free process, while enabling their integration into flexible substrates, as we have previously reported [13]. The suitability of these transfer-free graphene electrodes for multimodal applications was successfully demonstrated, showing no photo-induced artifacts and compatibility with MRI, further highlighting their potential for advanced neural interfacing technologies.

In this study, we describe lithography-compatible transfer-free multilayer graphene MEAs directly fabricated on transparent wafer substrates, with electrode sizes as small as 10-50 µm in diameter, comparable to or smaller than state-of-the-art transparent microelectrodes [23], [24], [25], [26], for in vitro multimodal platforms. We show that transfer-free multilayer graphene achieved impedance values lower than those of previously reported CVD monolayer or few-layer graphene electrodes. This could be attributed to an increase in the intrinsic graphene quantum capacitance with the number of layers, in accordance with previous work [20], [27]. Most importantly, the proposed MEAs enabled electrophysiological recordings of spiking activity from brain slices with high signal quality. Overall, the obtained results show that transfer-free multilayer graphene can be effectively integrated into in vitro MEA platforms and offers a great promise for advanced multimodal neural interfaces.

## 2. Results and Discussion

### 2.1. Fabrication and characterization of transfer-free graphene MEAs

To enable multimodal recordings, we designed and fabricated graphene-based MEAs on transparent fused silica substrates. Several electrode arrays with varying electrode sizes, numbers of electrodes, and pitches were designed to meet diverse experimental requirements and explore their technological limits. The primary designs included a 60-electrode MEA with 50 µm diameter electrodes arranged in an 8×8 square grid with a 200 µm pitch, and a 32-electrode MEA with 30 µm diameter electrodes arranged in a 6×6 grid with a 100 µm pitch. Additionally, a MEA with various electrode diameters, in the range of 10 to 500 µm, was included for a more direct and efficient characterization of the electrodes’ performance with respect to their sizes. The graphene layer, which defined the microelectrodes and inner tracks, was confined to the designated field of view to ensure optical transparency for neural imaging. Outside this region, a highly conductive and biocompatible metal, Ti/Au, was used to form the outer tracks and contact pads (see **Figure** 1a). A custom-designed printed circuit board (PCB) electrically interconnected to the MEA pads ensures efficient interfacing with commercially available electrophysiology recording systems, as shown in Figure 1d and Figure S4, Supplementary Information.

**Figure 1.**
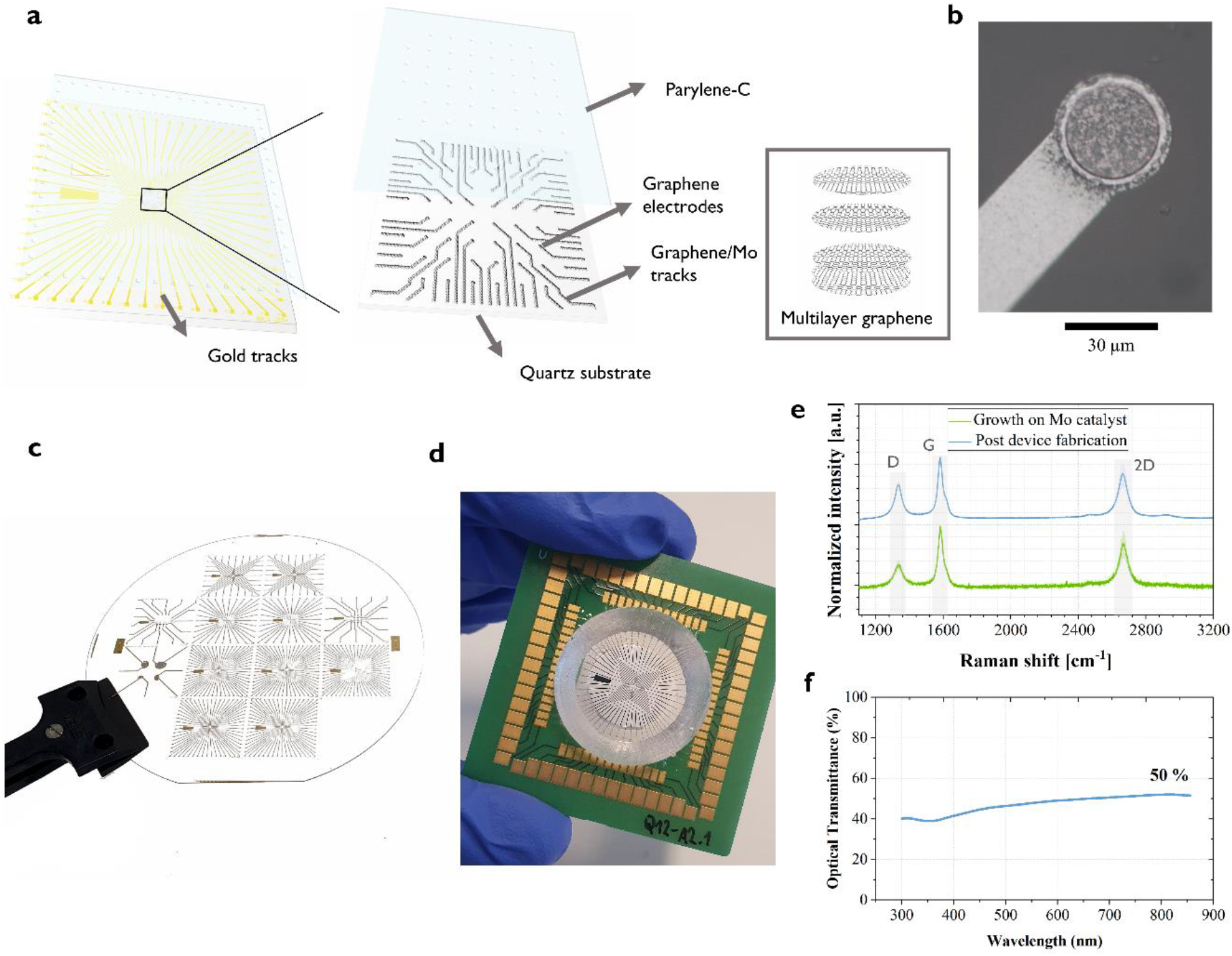
Design and fabrication of transfer-free graphene microelectrode arrays. a. Schematic of the graphene MEA device architecture and material layers. b. Optical image of a multilayer graphene electrode directly grown on the patterned molybdenum layer. c. Photograph of a 4 in. fused silica wafer containing 12 MEAs with different array designs. d. Graphene MEA device used for the in vitro recording experiments. e. Averaged Raman spectra (n = 5) of transfer-free multilayer graphene obtained immediately after chemical vapor deposition (CVD) growth on the Mo catalyst and after completing the MEA microfabrication process, confirming the structural integrity and quality of the graphene throughout the fabrication steps. Additional Raman spectra at different stages of the fabrication process are provided in the Supporting Information (SI). f. Optical transmittance measurements of transferred graphene sheets, with the contribution from the glass substrate subtracted.

The microfabrication process for transfer-free graphene MEAs was initiated by the deposition and patterning of a molybdenum (Mo) layer on a fused silica wafer. Mo acts as the catalyst layer for the subsequent multilayer graphene growth via chemical vapor deposition (CVD) [28]. To ensure the integrity of the graphene recording sites during the subsequent processing steps, a protective Ti/Al layer was deposited and patterned using a lift-off technique. We also attempted deposition via sputtering followed by wet etching. However, this often resulted in non-uniform etching process, leading to overetched structures, severely compromising the final MEA device functionality. Ti/Au tracks and contact pads were then formed via evaporation and lift-off. The devices were encapsulated with a 1–2 µm-thick parylene-C film, deposited via CVD, and patterned to expose the recording sites and contact pads. The fabrication process was completed with the removal of the Ti/Al protective layer and underlying Mo layer, leaving only graphene exposed on the electrodes. An optical image of the multilayer graphene electrode is shown in Figure 1b. Additional details on the microfabrication process are provided in the Experimental Section, and a schematic representation is provided in Figure S1 (Supporting Information).

Raman spectroscopy revealed the multilayer nature of the CVD grown graphene films. The characteristic G, 2D and D peaks of graphene (∼1582, 2660, 1335 cm ^-2^, respectively) were observed, with an I_2D_/I_G_ ratio lower than 1, confirming the presence of the multilayer graphene (see Figure 1e). Additionally, the single-peak nature of the 2D peak strongly suggested that the graphene layers were turbostratic, as they did not exhibit the complex splitting of more ordered stacking [29]. Turbostratic graphene refers to a type of multilayer graphene where the layers are rotationally misaligned, leading to weaker interlayer interactions and electronic decoupling [30]. The I_D_/I_G_ ratio, indicative of the amount of defects, was low, I_D_/I_G_ = 0.35, right after graphene growth on Mo catalyst layer on a quartz substrate. The number of defects slightly increased as the graphene layer was post-processed. Additional Raman spectra at different stages of the fabrication process are provided in Figure S3 (Supporting Information).

Optical transparency within the field of view of the microelectrode array is a critical requirement for enabling multimodal studies. To address this, our design employed multilayer graphene grown on a Mo catalyst layer to define the inner tracks within the field of view rather than traditional gold tracks. Gold, while highly conductive, exhibits photoelectric effects that can interfere with optical modalities, whereas multilayer graphene does not suffer from such limitations [13]. Optical transmittance analysis of the multilayer graphene in this work revealed transparency levels of approximately 50% across the entire spectrum, as shown in Figure 1f. A transmittance of 47.5% was obtained at a wavelength of 550 nm, corresponding to an absorbance of 52.5%. Based on the typical absorption of ∼2.3% per graphene layer [31], this corresponds to roughly 20–25 layers, assuming a linear approximation. While such approximations are less accurate at higher layer counts due to interlayer optical effects, they provide a useful estimate of the film’s thickness. Though this transparency is lower than that of monolayer graphene or ITO-based systems (typically >80%), it remains within a workable range for in vitro imaging modalities, and can be further optimized by tuning the CVD growth process. Notably, while the Mo layer was completely etched away from the electrode sites, it remained beneath the inner tracks, which could slightly impact the overall transparency within the field of view. Ongoing efforts are focused on developing methods to completely remove the Mo layer after graphene growth, thereby further enhancing the optical performance for multimodal applications.

The electrochemical properties of the transfer-free MEAs are detailed in Figure 2. The impedance dependency on the electrode size was studied by measuring the electrical impedance response of graphene electrodes of different sizes, ranging from 10 to 500 μm in diameter. Figure 2b shows the average impedance magnitude at 1 kHz plotted against the electrode diameter. The impedance magnitude decreased with increasing electrode size, as expected, following an inverse relationship. The corresponding numerical values are summarized in Supporting Information, Table S2.

**Figure 2.**
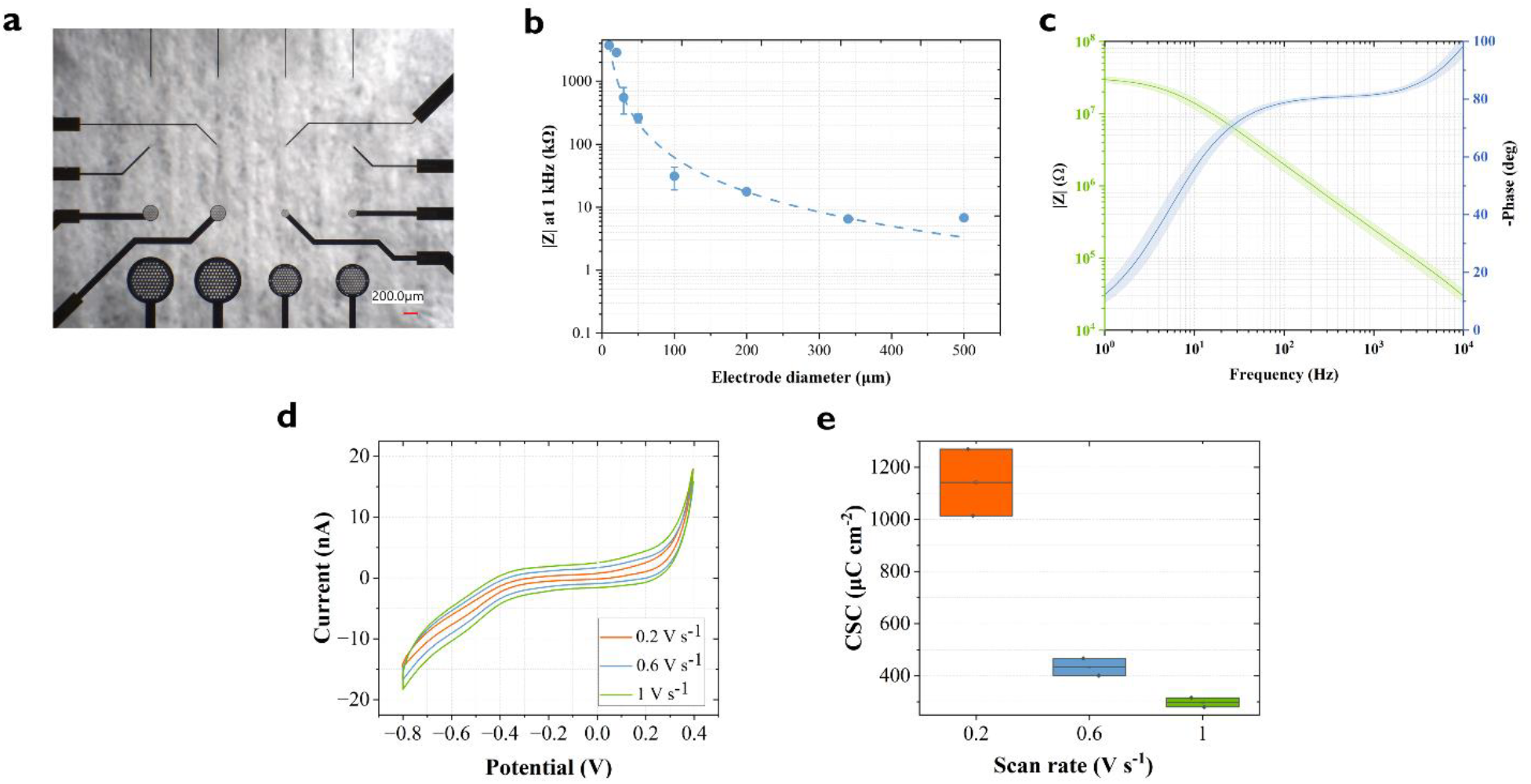
Electrochemical properties of graphene-based MEA electrodes. a. Optical image of MEA device featuring electrodes of different sizes. b. Electrochemical impedance spectroscopy (EIS) magnitude at 1 kHz of various electrode sizes, ranging from 10 to 500 µm. The dotted line represents the power fitting function of the data points. Circles represent the mean impedance values, with error bars indicating the standard deviation for electrode sizes with 10 measurements (30, 50 and 100 µm). For electrode sizes with only two measurements (10, 20, 200, 340, and 500 µm), the error bars were omitted because of insufficient data. c. Averaged impedance spectra (n = 10), including magnitude and phase, for 50 µm diameter electrodes. Shaded regions show standard deviation. d. Cyclic voltammetry (CV) curves for 50 µm diameter electrodes at scan rates of 0.2, 0.6 and 1 V s^-1^. e. Total CSC values extracted from the CV curves depicted in d.

Impedance spectra (*n = 10*), including magnitude and phase, are featured in Figure 2c for electrodes with a diameter of 50 µm. The average impedance magnitude at 1 kHz was 263.6 ± 44.0 kΩ (4.81 ± 0.80 Ω·cm^2^). The EIS magnitude follows the typical impedance behavior of neural recording electrodes, with an inverse relationship between the impedance and frequency. However, the EIS phase exhibits an unexpected tendency toward 0° at lower frequencies, which suggests the presence of a resistive component overshadowing the capacitive behavior. A plausible explanation for this deviation is the presence of Faradaic processes (redox reactions), potentially arising from Mo located beneath the graphene tracks. Although not in direct contact with the electrode surface, this underlying Mo may still interact with the electrolyte through exposed edges or defects in the graphene, thereby participating in electrochemical reactions. Mo is electrochemically active and can participate in redox reactions [32]. Although this does not impede the device’s ability to record high quality electrophysiological signals, complete Mo elimination is an important subject of this work and further investigation will be followed to confirm this hypothesis.

We then studied the reversibility of the electrochemical reactions on the electrodes and the amount of charge that can be stored reversibly, i.e., charge storage capacity (CSC), through cyclic voltammetry (CV). The CV curves for 50 µm diameter electrodes at scan rates of 0.2, 0.6 and 1 V s^-1^ are shown in Figure 2d. With the increase of the scan rate, the diffusion layer at the electrode-electrolyte interface becomes thinner, resulting in greater current densities. The water window was maintained between -0.8 V and 0.4 V to prevent undesirable electrochemical reactions. The CSC for each electrode was calculated by integrating the area under the CV curve within this water window and normalizing it to the electrode area. The average CSC values of the fabricated electrodes are shown in Figure 2e.

### 2.2. Electrophysiological recordings of neuronal activity from brain slices

The in vitro performance of transfer-free multilayer graphene MEAs, with 30 and 50 µm-diameter electrodes, was evaluated by recording neural activity from acute mouse cerebellar brain slices. The cerebellum was selected as the model system because of its well-characterized spontaneous spiking activity patterns, which are dominated by the firing of Purkinje cells. These cells exhibit regular and rhythmic spikes at a relatively constant and high frequency of around 30 to 50 Hz [34]. The preparation of cerebellar slice culture is described in the Experimental Section. The tissue slices were carefully positioned on the MEA surface and secured using a slice anchor with parallel nylon fibers to ensure close contact (see Figure 3b). Artificial cerebrospinal fluid (aCSF) was manually perfused over the slice to maintain tissue viability. We developed a custom-designed interface to facilitate the connection between the graphene MEAs and the Intan recording system (Intan Technologies, Los Angeles, CA). Further information regarding the acquisition setup is provided in the Experimental Section.

**Figure 3.**
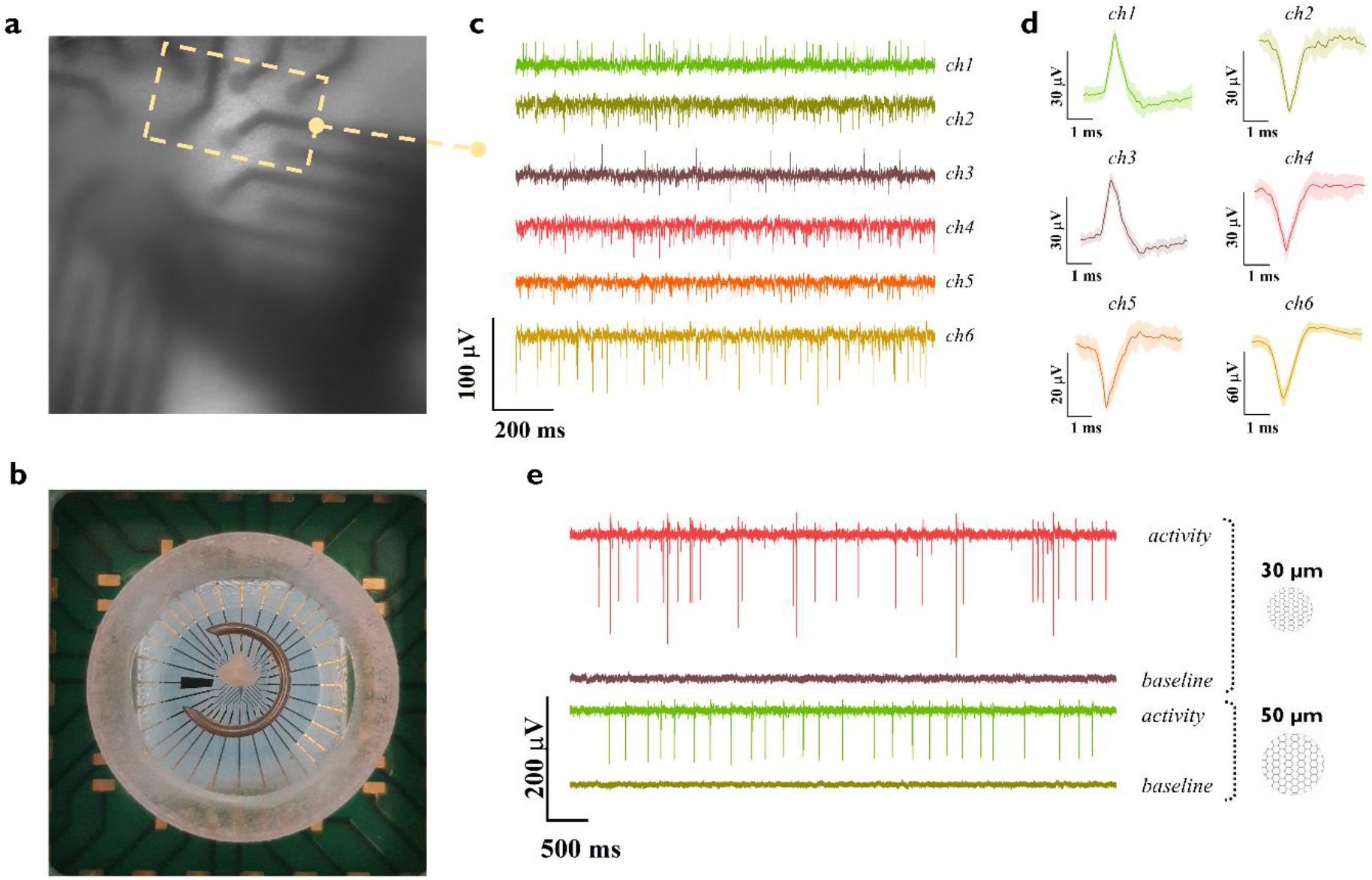
Electrophysiological recordings of neuronal activity from cerebellar brain slices. a. High-resolution optical image of the brain slice positioned over the MEA, captured with an epifluorescence microscope. b. Photograph (Top view) of a mouse cerebellar brain slice mounted on a graphene-based MEA device. To ensure close contact between the slice and the MEA, a slice anchor with parallel nylon fibers was gently pressed onto the tissue. c. Neural activity recorded by graphene MEA electrodes with diameter of 50 µm, showing the activity of six distinct channels. d. Averaged detected spike waveforms corresponding to the recording in c. Shaded regions represent standard deviation. e. Baseline and neural activity was recorded from graphene MEA devices with electrode diameters of 30 and 50 µm.

Prior to recording from a slice, a baseline signal was acquired for several minutes to assess the noise level of the system. The recorded filtered signals exhibited a root mean square (rms) noise of 2–3 µV, with smaller electrodes showing a higher noise floor. Spontaneous extracellular spiking activity of neuronal cells within the cerebellar slice was successfully detected, with spike amplitudes ranging from 6 to 300 µV across 30- and 50-µm diameter electrodes (Figure 3c–e). The amplitude variation can be attributed to several factors: (1) electrode size and spatial averaging—larger electrodes capture signals averaged over a broader area, potentially diluting contributions from individual neurons, whereas smaller electrodes resolve higher-amplitude signals from localized sources [35], [36]; and (2) neuron-electrode proximity—tight coupling between the tissue and electrode surface enhances signal amplitude. Additionally, variations in spike amplitude were observed within individual electrode recordings, indicating that multiple nearby neurons were being captured simultaneously, with each contributing spikes of different magnitudes. Moreover, different neurons are expected to have different spike waveforms [37].

The quality of the recorded electrophysiological signals was quantified by the signal-to-noise ratio (SNR), following the methodology described in the Experimental Section. The average SNR values of the filtered signals were found to be 19.38 ± 7.21 dB for the 30 μm diameter graphene electrodes and 20.58 ± 6.10 dB for the 50 μm electrodes. The distribution of the obtained SNR values is depicted in Figure 4c. Despite the difference in electrode size, the SNR values were found to be similar. This could be attributed to the inherent properties of smaller electrodes: while they are capable of detecting neural signals with higher amplitude, they are also more susceptible to increased levels of noise. It should be noted that the comparison of the SNR with previously reported values in the literature for transparent recording electrode arrays is challenging, as there is no standardized approach to compute the SNR. While some reports use peak-to-peak spike amplitudes relative to the baseline noise standard deviation, others employ RMS calculations or different filtering approaches. Moreover, electrophysiological signal characteristics vary substantially depending on the experimental paradigm, whether drug-induced, evoked, or spontaneous activity. Despite this, our results demonstrate the capability of transfer-free graphene MEAs to reliably detect extracellular neural activity. The achieved SNR values, particularly considering that the recordings were obtained from spontaneous activity, validate the potential of the technology for neurophysiological research applications.

**Figure 4.**
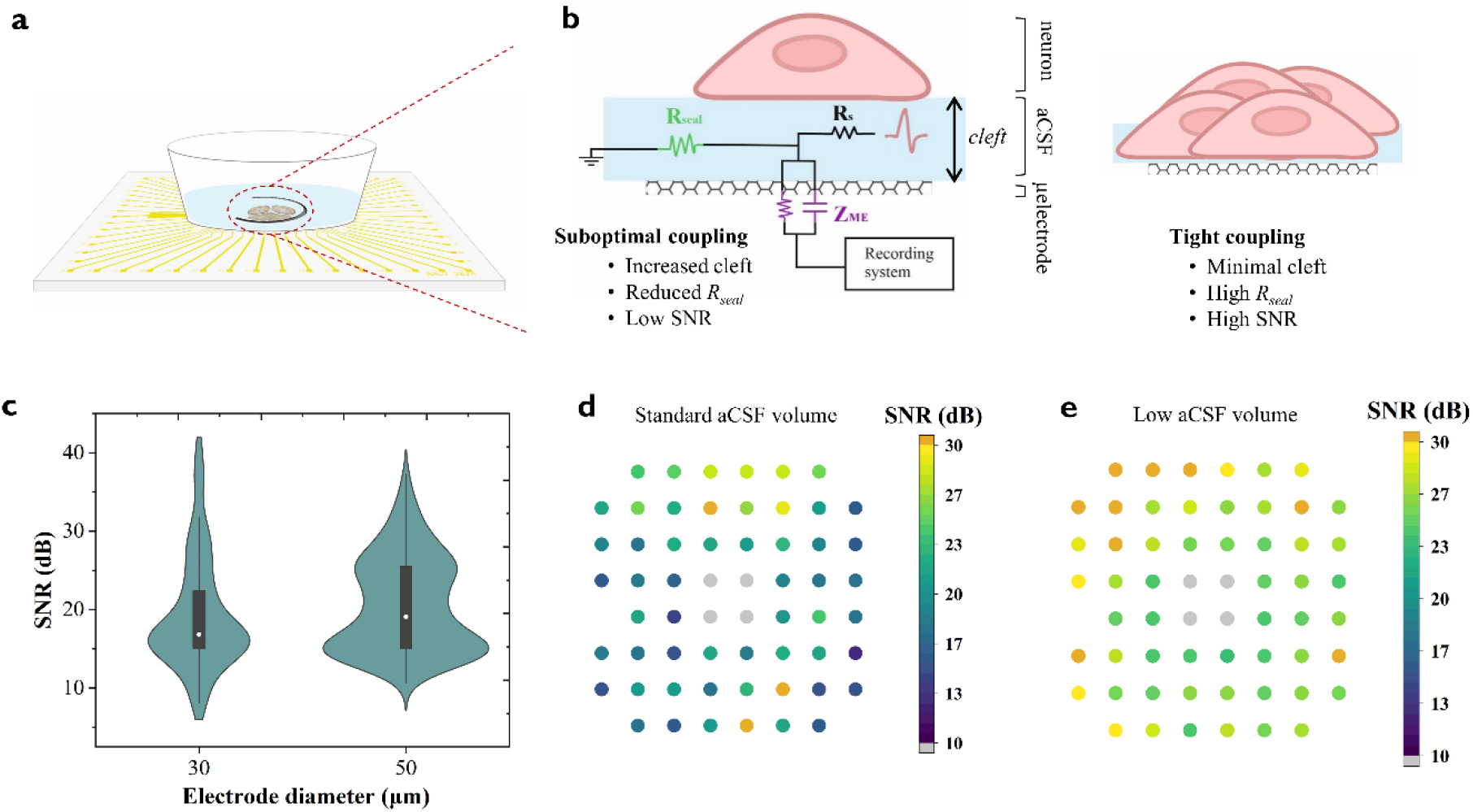
The electrical coupling between the neuronal cells and the microelectrodes. a. Schematic representation of the mouse cerebellar brain slice mounted on the fabricated MEA device and perfused with artificial cerebrospinal fluid (aCSF) to maintain tissue viability. b. Simplified contact model of the neuron-electrode coupling, adapted from Spira and Hai [ref2]. The recorded signal quality largely depends on the electrode impedance (ZME) and on the seal resistance (Rseal). c. Obtained signal-to-noise ratio (SNR) values per each tested electrode size and considering various activity recording sessions with different brain slices. d. Colormap representation of SNR values obtained with 50 µm-diameter electrodes overlaid on the MEA. Electrodes marked in grey indicate damaged ones, defined by an impedance magnitude greater than 1 MΩ at 1 kHz. e. Colormap representation of SNR values, similar to d, for recordings performed with a low aCSF volume.

Notably, the signal quality was found to be predominantly governed by the electrode impedance and neuron-electrode coupling. Electrode impedance is traditionally regarded as a primary benchmark for neural recording performance. However, in practice, electrode impedance only needs to be sufficiently low such that its thermal noise and voltage-divider effects do not dominate over the intrinsic characteristics of the recording electronics. Beyond this threshold, further reductions in impedance yield diminishing returns in signal quality. Compared to CVD monolayer or few-layer graphene, the transfer-free multilayer graphene microelectrodes presented here exhibit reduced impedance due to their increased effective surface area, while also eliminating fabrication variability and defects associated with the graphene transfer process. Other transparent electrode materials, such as PEDOT:PSS, indium tin oxide (ITO), and MXenes, have been explored for multimodal neural interfacing [23], [24], [33]. Among these, PEDOT:PSS has demonstrated exceptionally low impedance owing to its volumetric capacitive behavior; however, its long-term chemical and mechanical stability remain significant challenges, particularly in complex biological environments. Although the impedance of our MLG electrodes remains higher than that of PEDOT:PSS-coated devices, it is comparable to or lower than values reported for ITO and MXene-based electrodes. More importantly, we demonstrate that these MLG electrodes achieve high-fidelity recordings of spontaneous extracellular activity in acute brain slices, with low noise, spike amplitudes up to 300 µV, and signal-to-noise ratios reaching 30–40 dB—despite their higher impedance.

The coupling quality can be characterized by the seal resistance (R_seal_) [38], [39], [40]. This key parameter represents the electrical resistance to current leakage through the extracellular cleft between the neuronal membrane and electrode surface (see Figure 4b). R_seal_ exhibits a strong inverse relationship with the physical neuron-electrode distance. Our experiments demonstrated that optimal recordings (SNR > 30 dB) were achieved when the electrodes were in close proximity to active neurons and in tight contact with the tissue, thereby minimizing the cleft distance. However, perfusion with aCSF led to gradual tissue detachment from the electrode surface, increasing cleft dimensions and thereby, reducing R_seal_, from the expansion of the fluid-filled gap between cells and electrodes. While various noise sources contribute to signal degradation, including thermal noise associated with electrode impedance, biological noise from neural tissue, and electronic noise from recording instrumentation [41], in our in vitro recordings, we found that the dominant factor determining signal quality was the quality of neuron–electrode coupling, rather than absolute impedance alone. These findings underscore that achieving high quality neural recordings requires optimization of both electrode properties and neuron-electrode electrical coupling for in vitro experiments. In this context, we demonstrate that excellent neural signal acquisition is achievable without ultra-low impedance, as long as electrode design, tissue coupling, and recording chain are appropriately matched. Our transfer-free multilayer graphene (MLG) electrodes, reliably capture spontaneous extracellular activity with high signal-to-noise ratios. This establishes transfer-free MLG as a robust and high-performance platform for transparent neural interfacing.

## 3. Conclusion

In this study, we developed transfer-free graphene MEAs for interfacing with neural tissue and demonstrated their capability for electrophysiological recording. Our transfer-free fabrication approach represents a significant advancement in graphene-based neural technology, addressing the reliability and scalability challenges associated with state-of-the-art CVD graphene wet transfer methods. We showed that transfer-free graphene can be miniaturized to the micrometer scale, down to 10 µm in diameter, and integrated into transparent in vitro substrates, while preserving the integrity of the graphene film. [13]

The electrochemical characterization revealed low impedance values (263.6 ± 44.0 kΩ at 1 kHz for 50 μm-diameter electrodes), which enabled the detection of spontaneous neural activity from acute cerebellar brain slices. Our experiments revealed that the neuron-electrode coupling strength was the main limiting factor for signal fidelity. Optimizing the coupling between the neural tissue and the electrode surface in in vitro platforms remains a critical aspect for achieving high-quality neural recordings. Despite this limitation, several electrodes recorded neural signals with exceptional signal quality (SNR > 30 dB), demonstrating the potential of this technology for efficient neural interfacing.

In addition to advancing the understanding of electrophysiological recording capabilities, this study emphasizes the broader potential of transfer-free graphene MEAs for multimodal neural interfacing. Their compatibility with optical imaging is ensured by restricting the field of view to non-photoelectric materials (e.g., graphene grown on a Mo layer and transparent substrates), paving the way towards fully transparent electrode arrays. Furthermore, prior work [13] has demonstrated that transfer-free graphene electrodes exhibit no photo-induced artifacts and remain MRI-compatible, highlighting their suitability for multimodal applications. These properties position transfer-free graphene MEAs as a promising platform for integrating electrical and optical modalities, providing high spatiotemporal resolution for comprehensive neural network analysis.

Overall, this work demonstrates that transfer-free graphene MEAs offer an innovative approach to neural interfacing by overcoming current fabrication limitations and providing reliable, high-quality electrophysiological recordings. The integration of electrical and optical modalities within a single transparent platform opens new avenues for advancing neuroscientific research. Future research directions should focus on optimizing neuron-electrode coupling, exploring bidirectional functionalities and translating this technology to in vivo settings, ultimately contributing to the next-generation of multimodal neural interfaces for a deeper understanding of complex brain functions and associated disorders.

## 4. Experimental Section

### 4.1. Device fabrication and assembly

Fused silica quartz wafers were used as transparent starting material due to their ability to withstand the high temperatures required for graphene growth, their superior resistance to graphene delamination compared to sapphire substrates and their ease of dicing in comparison to sapphire [42]. The microfabrication process of the graphene MEAs is illustrated in Figure S1 (Supporting Information).

The fabrication started with a 50 nm molybdenum (Mo) layer deposition at 50ºC in the frontside of the wafer via sputtering. Mo serves as the catalyst layer for the subsequent graphene growth. Next, a temporary 50 nm Ti layer was sputtered at 50ºC at the backside for the wafer to be optically detected in the metal etcher equipment. After both sputtering processes, standard lithography steps defined the pattern in the Mo layer using a positive photoresist (SPR3012, Shipley Megaposit) (Figure S1b). The lithography process required adaptation, such as extended bake times, to account for the lower thermal conductivity of quartz. The defined pattern was then transferred by plasma etching of the Mo layer in an ICP-RIE etcher. The etching was performed at 40ºC with no RF platen power, 500 W ICP power, 5 mTorr pressure, and SF_6_ gas at 25 sccm flow. Following this, the ion bombarded photoresist was stripped using a bath with NI555 solvent solution. Ultrasonication at 80 kHz and elevated temperatures (50-60ºC) were employed to accelerate the process. The Ti layer in the backside was then fully wet etched using HF 0.55% bath to avoid any contributions from it during the graphene growth process. Graphene was then selectively grown on the Mo structures. The growth was done via chemical vapor deposition (CVD) process at 935ºC with 960, 40 and 25 sccm of Ar, H_2_ and CH_4_ gases for 20 min at 25 mbar.

A Ti/Al metal layer protected the graphene electrodes throughout the fabrication process and in particular, during the plasma etching of the Parylene-C encapsulation layer. After the lithography using a negative photoresist (nLOF-2020, Microchemicals GmbH, Germany), a stack composed of 10 nm Ti and 100 nm pure Al was evaporated. For the patterning of the metal layer via lift-off, NI555 stripper was used to remove the photoresist and along with the evaporated metal on top, leaving the protective Al structures on the electrodes revealed (Figure S1d).

Before preparing the wafers for Au evaporation to form the outer MEA tracks and contact pads, a 200 nm Ti layer was evaporated in the backside. The Ti layer enabled the electrostatic clamping in the ICP-RIE plasma etcher process step to come. The Au layer was e-beam evaporated and defined via lift-off process by using a negative photoresist (nLOF-2020, Microchemicals GmbH, Germany) (Figure S1e). The photoresist was stripped with NI555 solvent solution overnight.

The wafers were then coated with a 1-2 μm parylene-C layer (Figure S1f), which was CVD deposited at room temperature. Parylene-C served as the insulating coating of the MEAs. 3-(trimethoxysilyl)propyl methacrylate (A-174 Silane) was used as an adhesion promoter for parylene on the substrate. A patterned thick positive photoresist coating (AZ10XT, Microchemicals GmbH, Germany) was used as the masking layer to expose electrodes and contact pads in the parylene layer. A plasma etching step created the openings in the parylene-C, landing on the Al protective layer. O_2_ and SF_6_ gases were used at 185 sccm and 15 sccm flows respectively, and with 40 W LF power. The photoresist mask was removed with acetone followed by IPA. Additional cleaning with NI555 was performed to ensure that the ion bombarded photoresist was completely removed.

Prior to dicing, the whole wafer surface was coated with a thick positive photoresist layer (AZ3027, Microchemicals GmbH, Germany) to protect it from the dicing procedure. The quartz wafers were then diced with a dedicated diamond blade. The microfabricated dies were cleaned in acetone followed by IPA. Thereafter, the Al protective layer on the electrodes as well as the backside Ti layer were fully etched in 0.55% HF solution. Finally, Mo was wet etched by gently covering the electrode area with 31% H_2_O_2_ forming a puddle (Figure S1i). An etching time of 3 min was sufficient to remove almost completely the Mo layer underneath without causing graphene detachment. The microfabricated dies were left in vacuum overnight at 80ºC to remove any moisture and ensure a better adhesion in between layers.

The microfabricated dies were attached and electrically interconnected to a custom-designed PCB via wire bonding to facilitate electrode characterization and ensure efficient interfacing with commercially available electrophysiology recording systems. The design of the PCB, including the pads’ placement, size, and pitch, was specifically optimized to be compatible with Multichannel Systems MEAs featuring 60 electrodes. This configuration also ensured compatibility with other commercial electrophysiology systems, such as the MZ60 MicroElectrode Array interface from TDT.

To contain phosphate-buffered saline (PBS) or culture medium during experiments, inverted cone-shaped wells were designed and fabricated using an Asiga3D printer. A biocompatible resin, Detax Freeprint Ortho, was selected for 3D printing the wells to minimize the risk of cytotoxicity or other adverse biological responses when in contact with cells or tissue slice cultures [43]. The 3D-printed wells were securely adhered to the surface of the MEA device using the same non-conductive epoxy (EPO-TEK 301-2FL), which also served to cover the wirebonds, providing additional mechanical protection against physical damage during handling and use. Figure 1d shows a completely fabricated and assembled graphene MEA device.

### 4.2. Device characterization

#### Raman analysis

Raman spectroscopy was employed to confirm the presence of graphene, and extract information about the type of graphene, quality and amount of defects present in the grown layer. Raman spectra were obtained using a Renishaw inVia Raman system with a laser of 633 nm. The spectrum was acquired with 50% laser power and 20 s of exposure time to achieve an adequate signal-to-noise ratio. Several point measurements from each sample were taken, and then analyzed with a custom Matlab script.

#### Optical transmission

To investigate the level of transparency of the final device, optical transmittance measurements were conducted. For these measurements, a cm-scale graphene layer was transferred onto a glass substrate. PerkinElmer Lambda 1050+ UV/VIS/NIR spectrometer was used to evaluate the optical transmittance of graphene samples. Transmittance data over a wide range of wavelengths, from 300 nm to 860 nm, was obtained. Measurements were taken for graphene and glass samples, allowing the isolation of the glass’s contribution.

#### Electrochemical characterization

Standard electrochemical measurements were conducted to characterize microelectrodes’ performance. Electrochemical impedance spectroscopy (EIS) and cyclic voltammetry (CV) were performed in a three-electrode setup and using a phosphate buffered saline (PBS), PBS 1X pH 7.4, as the electrolyte solution. The three-electrode setup consisted of: a Pt electrode (3 mm diameter (BASI Inc.)) serving as the counter electrode (CE), a leakless miniature silver/silver chloride (Ag/AgCl) (eDAQ) as reference electrode (RE), and the fabricated graphene microelectrodes as the working electrodes (WE). All the electrodes were connected to a potentiostat (Autolab PGSTAT302N) and kept inside a Faraday cage. For the EIS, a 10 mV sine-wave voltage was applied between the WE and RE, and the current between the WE and CE was measured. The impedance magnitude and phase over a range of frequencies (from 0.1 Hz to 100 kHz) was recorded. For the CV, the water window, defined as the potential range where water remains stable and avoids electrolysis, was set from -0.8 to 0.4 V for the 50 µm-diameter electrodes. CV measurements at several scan rates were taken: 0.2, 0.6 and 1 V s^-1^.

### 4.3. MEA acquisition setup

Neural activity was acquired and amplified using Intan RHD amplifier chips (RHD2132) and digitized by the Intan RHD USB Interface Board (Intan Technologies, Los Angeles, CA). To facilitate the connection between the MEA devices and the Intan recording system, a custom-designed interface was developed. This setup, illustrated in Figure S4, included a custom-printed circuit board (PCB) that interfaced the MEA pads with Omnetics connectors, which were linked to two 32-channel Intan recording headstages (containing RHD2132 chips). The RHD USB Interface Board was connected to a computer running RHX Data Acquisition Software, enabling real-time data collection and analysis.

The custom PCB was designed to support up to 60 channels (59 recording electrodes and one internal reference electrode), ensuring compatibility with both the fabricated graphene-based MEAs and standard 60-channel commercial MEAs from Multichannel Systems GmbH. Spring-loaded pins, soldered to the custom PCB, ensured reliable electrical contact with the MEA pads. To maintain stable mechanical coupling between the MEA and the custom PCB, laser-cut components were fabricated from 2 mm-thick plexiglass sheets. These components were designed to securely host the MEA while maintaining proper alignment between the MEA pads and the spring-loaded pins on the PCB. By applying pressure through screws mounted on the assembly, the spring-loaded pins were pressed against the MEA pads, ensuring reliable and consistent electrical contact (Figure S4c).

### 4.4. Acute cerebellar slice preparation

All animal experiments were approved by the national Central Commissie Dierproeven and the institutional animal welfare committee of Erasmus MC. A male C56/BL6 mice of 3 months old, bred at the in-house animal facility of Erasmus MC, were decapitated under isoflurane anesthesia. Subsequently, the cerebellum was removed and transferred into ice-cold slicing medium containing (in mM): 240 sucrose, 5 KCl, 1.25 Na2HPO4, 2 MgSO4, 1 CaCl2, 26 NaHCO3, and 10 D-glucose, bubbled with 95% O2 and 5% CO2. Parasagittal slices 250 μm thick of the cerebellar vermis were cut using a Leica vibratome (VT1000S, Nussloch, Germany) and kept in artificial cerebrospinal fluid (aCSF) containing (in mM): 124 NaCl, 5 KCl, 1.25 Na2HPO4, 2 MgSO4, 2 CaCl2, 26 NaHCO3, and 20 D-glucose, bubbled with 95% O2 and 5% CO2 for >1 h at 34 ºC before the experiments started.

### 4.5. Electrophysiological recordings

Extracellular activity from cerebellar brain slices was acquired at a 20 kHz sampling rate, using the MEA-to-Intan interface system described in section 4.3. The complete acquisition setup was placed inside a Faraday cage, with the ground pin of the Intan RHD USB Interface Board connected to it, establishing a common ground reference. ACSF was manually perfused over the slice to maintain tissue viability. To ensure close contact between the slice and the MEA, a slice anchor with parallel nylon fibers was gently pressed onto the tissue. Prior to recording from a slice, a baseline signal was acquired for several minutes to assess the noise level of the system.

### 4.6. Data analysis

Recordings were post-processed with a custom Matlab script using Signal Processing Toolbox. The data processing technique was based on previous work [37], [44]. The first step involved filtering the signals using a second-order IIR bandpass filter from 5 Hz to 1000 Hz, effectively isolating the frequency components relevant to local field potential (LFP) and spiking activity. Because IIR filters can largely distort the spike shapes due to phase nonlinearities, a zero-phase digital filtering was applied. After filtering, spikes were detected using an amplitude thresholding method based on the median absolute deviation (MAD) of the filtered signal. The detection threshold was initially set at 6 times the MAD. For the averaged detected spike waveforms depicted in Figure 3d, this threshold was readjusted by visual inspection to isolate higher amplitude spikes typically corresponding to neurons located closer to the recording electrode. This approach, while requiring manual intervention, provided a fast and straightforward method to minimize contributions from distant neurons, ensuring a more accurate representation of nearby neural activity. Additionally, to avoid detecting multiple spikes within the same action potential event, a refractory period of 2.5 ms was applied.

Finally, the signal-to-noise ratio (SNR) was computed by calculating the ratio of the maximum peak-to-peak amplitude of the neural spikes, *V*_*pp,signal*_, to the standard deviation of the noise in the baseline signal, *σ*_*noise*_, as shown in the expression below:

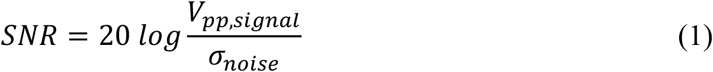

## Acknowledgements

The authors thank the Bioelectronics group at Delft University of Technology for their support and fruitful discussions. Special thanks go to Maria Camarena Perez for her insightful contributions to fabrication challenges. Authors thank Dr. Tiago Costa and Zu Yao Chang for their assistance with wafer dicing and wire bonding. This research used resources of the Else Kooi Laboratory (EKL) cleanroom facilities of Delft University of Technology, and the authors thank the EKL staff for their expert guidance throughout the microfabrication processes.

## Funding

The author(s) declare that financial support was received for the research, authorship, and/or publication of this article. This publication is part of the DBI2 project (024.005.022, Gravitation), which is financed by the Dutch Ministry of Education (OCW) via the Dutch Research Council (NWO).

## Conflict of Interest

The authors declare no conflict of interest.

## Notes

### Competing Interest Statement

The authors have declared no competing interest.

