## Supplementary material for "Transparent transfer-free multilayer graphene microelectrodes enable high quality recordings in brain slices": grMEA_SI_2025_final.pdf

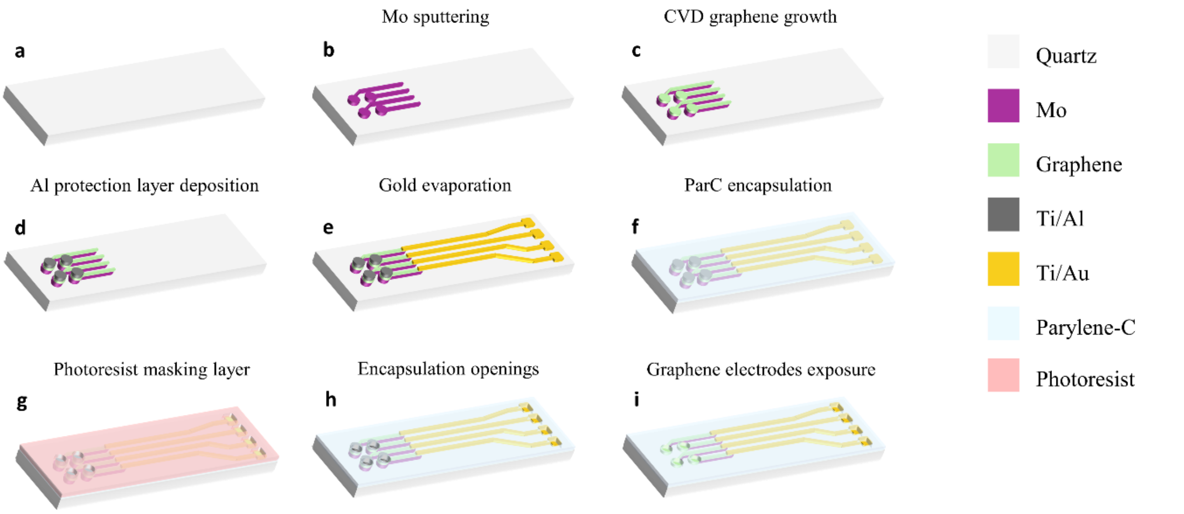


**Figure S1.** Wafer-scale transfer-free fabrication process steps of grMEA electrodes. a. Quartz wafer substrate, b. sputtering of 50 nm Mo layer and patterning, c. CVD multilayer graphene growth, d. Ti/Al deposition and patterning on the microelectrodes, and deposition of a 200 nm Ti layer in the backside (for the later electrostatic clamping in the AMS110 plasma etcher, in step h), e. Ti/Au (10/200 nm) evaporation and lift-off, f. deposition of 1 µm parylene-C encapsulation layer, g. photoresist mask deposition, h. parylene plasma etching, i. Ti/Al and Mo wet etching on the electrode openings.

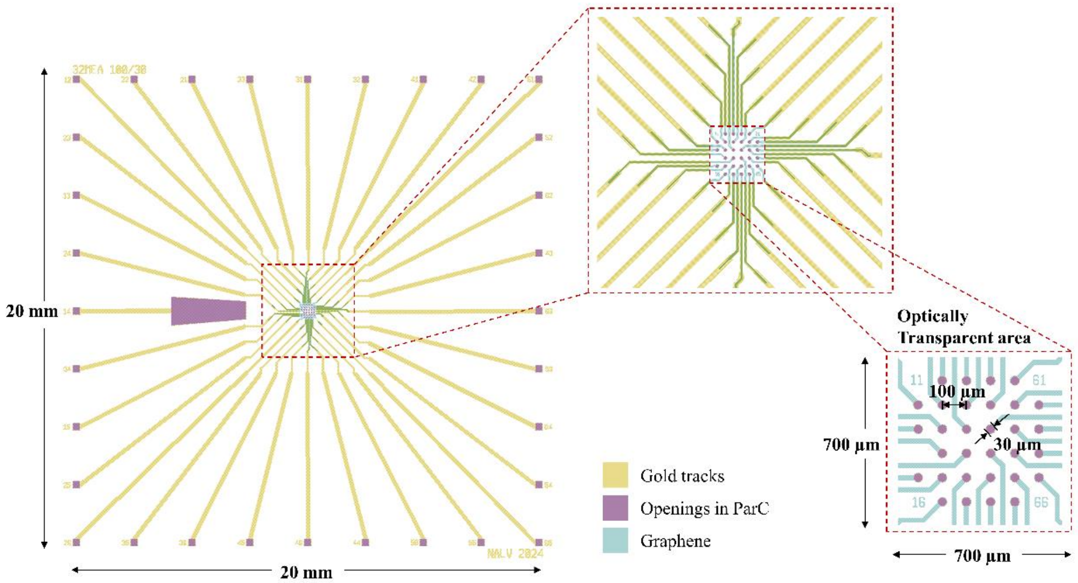


**Figure S2.** 32MEA100/30 electrode layout designed for the brain slice recording test. 31 working electrodes and one reference electrode are arranged in a 6×6 layout grid with electrode diameters of 30 µm and interelectrode distances of 100 µm. The electrode area, 700×700 µm, is designed to be optically transparent by forming the inner tracks with multilayer graphene grown on a temporary Mo catalyst layer.

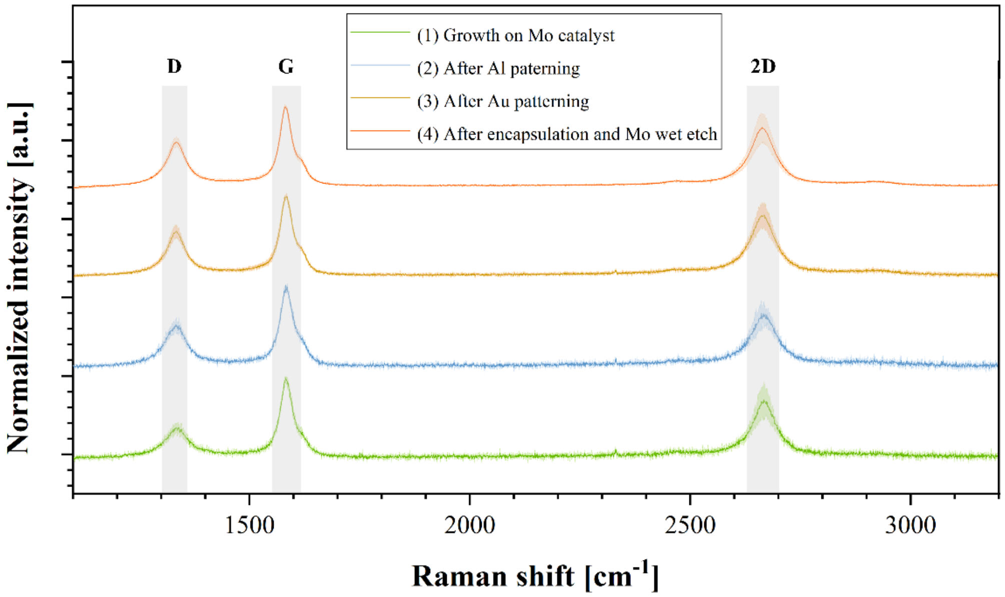


**Figure S3.** Raman spectra of the transfer-free multilayer graphene obtained at different stages of the microfabrication process: (1) Right after graphene grown on Mo catalyst on a quartz substrate, (2) After Al sacrificial protective layer deposition and patterning, (3) after Au tracks deposition and patterning, (4) after parylene C encapsulation, openings formation and Mo wet etching. Each graph line shows the average and standard deviation of 5-point measurements.

| *Process stage* | $I_D/I_G$ *ratio* | $I_{2D}/I_G$ *ratio* |
|---|---|---|
| Growth on Mo catalyst | $0.35 \pm 0.09$ | $0.72 \pm 0.25$ |
| After Al patterning | $0.49 \pm 0.11$ | $0.64 \pm 0.21$ |
| After Au patterning | $0.55 \pm 0.14$ | $0.78 \pm 0.19$ |
| After encapsulation and Mo etch | $0.50 \pm 0.08$ | $0.68 \pm 0.10$ |

**Table S1.** Raman ratios of transfer-free multilayer graphene obtained from the spectra at different stages of the process.

| | Impedance magnitude at $1kHz$ ($k\Omega$) | | | | | | | |
|---|---|---|---|---|---|---|---|---|
| Electrode diameter ($\mu m$) | 10* | 20* | 30 | 50 | 100 | 200* | 340* | 500* |
| Multilayer graphene | 3724.8 ± 286.1 | 2862.0 ± 501.2 | 550.2 ± 246.6 | 263.6 ± 44.0 | 31.1 ± 12.2 | 17.6 ± 0.1 | 6.5 ± 0.1 | 6.8 ± 1.4 |

**Table S2.** Impedance magnitude at 1 kHz (kΩ) of transfer-free multilayer graphene electrodes of various sizes, ranging from 10 to 500 μm in diameter. * The average and standard deviation are calculated from 2 measurements, limited by the number of electrodes available. Measurements for 30, 50 and 100 μm, instead, correspond to the average of 10 electrodes.

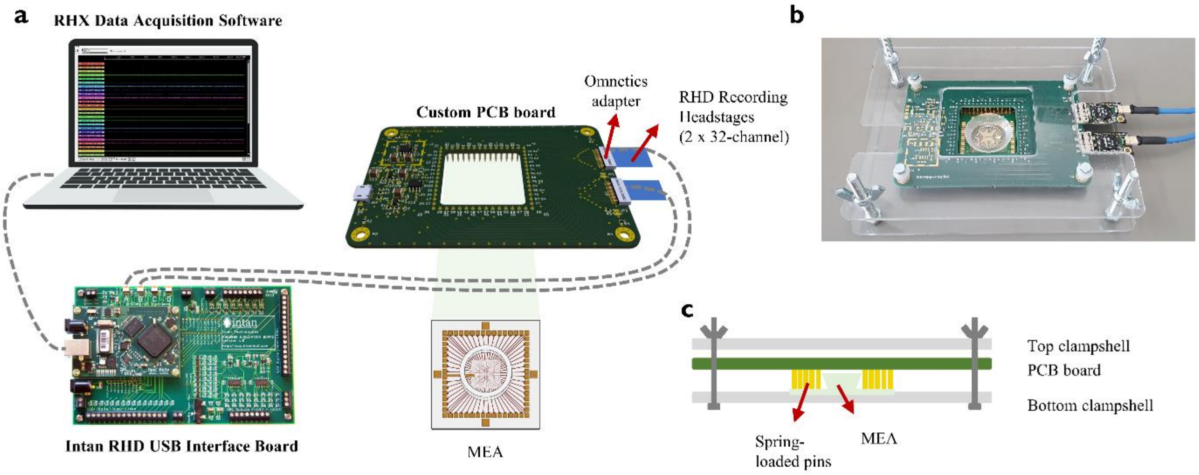


**Figure S4.** Overview of the MEA acquisition system setup. a. Schematic representation of the connections between the custom PCB board and the Intan recording system. The custom PCB board allows to couple the MEA electrodes, up to a total of 64, to the recording channels from Intan headstages. Spring-loaded pins, soldered to the custom PCB, effectively contact with the MEA pads and route these connections to the Omnetics adapter pads, and in turn to the Intan recording headstage pins. b. Image of the assembled MEA-to-Intan interface. c. Cross-sectional illustration of the MEA-to-Intan interface. The mechanical parts (top and bottom clampshells) ensure a good contact between the spring-loaded pins from the custom PCB and the MEA pads.
